# Neuro-Regenerative Behavior of Adipose-Derived Stem Cells in Aligned Collagen I Hydrogels

**DOI:** 10.1101/2023.05.12.539636

**Authors:** Mackenzie Lewis, Gabriel David, Danielle Jacobs, Alan E. Woessner, Patrick Kuczwara, Jin-Woo Kim, Kyle P. Quinn, Younghye Song

## Abstract

Peripheral nerve injuries persist as a major clinical issue facing the US population and can be caused by stretch, laceration, or crush injuries. Small nerve gaps are simple to treat, and the nerve stumps can be reattached with sutures. In longer nerve gaps, traditional treatment options consist of autografts, hollow nerve guidance conduits, and, more recently, manufactured fibrous scaffolds. These manufactured scaffolds often incorporate stem cells, growth factors, and/or extracellular matrix (ECM) proteins to better mimic the native environment but can have issues with homogenous cell distribution or uniformly oriented neurite outgrowth in scaffolds without fibrous alignment. Here, we utilize a custom device to fabricate collagen I hydrogels with aligned fibers and encapsulated adipose-derived mesenchymal stem cells (ASCs) for potential use as a peripheral nerve repair graft. Analysis of these scaffolds in vitro revealed heightened therapeutic secretome from ASCs, ECM deposition, and dorsal root ganglia (DRG) neurite outgrowth along the axis of fiber alignment. Our platform serves as an in vitro testbed platform to assess neuro-regenerative potential of ASCs in aligned collagen fiber scaffolds and may provide guidance on next-generation nerve repair scaffold design.

## Introduction

Peripheral nerve injury (PNI) remains a significant clinical challenge, with an average incidence of between 43 and 52 per one million people affected annually in the US alone^1,2^. While surgical coaptation of the severed ends of the nerves is possible in short- or no-gap injuries, long-gap injuries (20 mm or greater) require implantation of nerve repair grafts, including autografts, to avoid exerting excess tension on the remaining nerves and causing further damage. Nerve repair grafts are preferred over autografts to avoid additional surgery and corresponding sites of morbidity.

Current nerve repair implant strategies in the clinic are hollow nerve guidance conduits (NGCs) and decellularized nerve grafts^3^. While these strategies have been used to treat patients with nerve injuries, they do not provide universal solutions to nerve repair. Hollow NGCs are effective for short nerve gaps (less than 5 mm)^4^. On the other hand, decellularized nerve grafts such as Avance® nerve repair grafts have shown clinical success in longer gap repairs (up to 70 mm) because of the preservation of intraluminal basal lamina architecture. Nevertheless, decellularized nerve grafts possess a few limitations, including but not limited to 1) inability to tailor graft dimensions to individual patients, and 2) a need for migration of Schwann cells to facilitate axonal regeneration and axon myelination.

Research into recellularization of acellular nerve grafts has been conducted using Schwann cells or mesenchymal stem cells (MSCs), either undifferentiated or as Schwann-like cells, with abluminal coverage or endoneurial delivery using microinjectors^5–10^. While direct injection of Schwann cells is beneficial, clinical translation of Schwann cell transplantation remains challenging because of inherently low yield of autologous Schwann cells. While previous studies collectively show therapeutic benefits of MSC delivery in nerve-mimetic grafts, there still exists a need to develop grafts that allow an even distribution of MSCs across nerve grafts containing basal lamina-like topographical features. Microinjections of MSCs at four or more sites along the length of the grafts have been widely used, yet this approach relies on migration of injected cells for homogeneous distribution across the entire length of the nerves. This calls for a bottom-up approach in creating cellular nerve grafts containing nerve-mimetic topographical cues.

Aligned extracellular matrix (ECM) fibers, particularly collagen, have been shown to enhance nerve repair potentials^11–15^. This has been demonstrated in not only engineered nerve repair grafts containing aligned features, but also tissues such as tendons and muscles that contain aligned collagen fibers^16–18^. As such, there exist many different approaches to create aligned structures, including but not limited to uniaxial strain^19–21^; extracting longitudinal collagen fibers from inherently anisotropic tissues^22,23^; extruding precursor solutions to induce alignment of polymer fibers^13,24,25^; and creating aligned patterns via alignment of dissolvable magnetic beads^16,17^. Among these different approaches, applying uniaxial strain to collagen pre-gel solutions offers the most simplistic approach to align both collagen fibers and encapsulated cells throughout an entire three-dimensional (3D) hydrogel. Yet, it remains largely unclear how uniaxially aligned collagen I fibers modulate neuro-regenerative behavior of encapsulated adipose-derived MSCs (ASCs), and the potential of the combination as a highly effective nerve repair scaffold that promotes functional regeneration of severed nerves has not been well studied.

Here, we explore neuro-regenerative behavior of ASCs in 3D hydrogels containing uniaxially aligned collagen fibers. ASCs share many similarities with MSCs; however, one key advantage of ASCs over MSCs is their abundance and ease of isolation. These culture systems were created by uniaxial strain on ASC-encapsulated collagen pre-gel solutions cast onto a pre-stretched silicone mold. Our results show that uniaxial strain and compression of the gel from silicone release co-aligns collagen fibers and embedded ASCs. Furthermore, ASC-laden aligned gels exhibit mechanical properties that have been shown to promote Schwann cell migration, and aligned ASCs secrete significantly higher amounts of both neurotrophic and pro-angiogenic factors, deposit other neuro-regenerative ECM components such as fibronectin and laminin in an aligned fashion and promote dorsal root ganglia (DRG) neurite outgrowth along the aligned topographical cues. This effect may be in part modulated by YAP-mediated mechanotransduction. Overall, our study provides insight into neuro-regenerative behavior of ASCs in 3D aligned collagen I fiber scaffolds and further investigation may lead to the development of nerve repair scaffolds that contain nerve-mimetic intraluminal architecture and living depots of trophic factors.

## Materials and Methods

### Collagen I pregel formation

Collagen I was isolated from frozen rat tails (VWR RLRBT297) based on a previously established method^26^. Skin was first removed from the tail, and each vertebra was pulled to isolate the tendons. The isolated tendons were digested in 0.1% acetic acid (Millipore Sigma 1000631000) at 75 mL/g for 3 days at 4°C. The collagen digest was centrifuged in 40 mL volumes for 90 minutes at 8800 rpm and 4°C. The supernatant was frozen for 24 hours before being lyophilized (Labconco FreeZone 4.5) for 4 days. The resulting collagen I was digested in 0.1% acetic acid at a concentration of 10 mg/mL to create the pregel solution.

When used in hydrogel fabrication, 10% (v/v) 10X M199 (Sigma Aldrich M0650) were added to an aliquot of the pregel, followed by neutralization with 1M NaOH (Sigma Aldrich 415413). Phosphate buffered saline (PBS, VWR 97062) or cell suspension was then added to dilute to working concentration for each sample. These pregels were then incubated at 37°C to induce thermal gelation within their molds.

### hASC Culture

Human adipose-derived stem cells (hASCs, Lonza PT-5006) were cultured in ADSC Growth Medium Bulletkit (Lonza PT-3273 & PT-4503) containing 10% fetal bovine serum (FBS), 1% L-Glutamine, and 0.1% Gentamicin Sulfate-Amphotericin (GA-1000). All cultures were maintained at 5% CO_2_ and 37°C with media changes every 2-3 days. Cell passage p4-p5 were used for all cultures.

### Aligned Gels with ASCs

A custom stretching device was designed in SolidWorks and created with acrylic and steel (Figure 1a). PDMS sheets (SMI .010” NRV G/G 40D 12”X12”) were cut to the size of the silicone molds (Sigma Aldrich GBL664515) and plasma sealed using air plasma cleaner (Harrick Plasma) for >5 minutes (Figure 1f). The PDMS backed molds were clamped on both sides and placed on to the stretching device. The molds were then chemically functionalized to anchor collagen gels using a previously established method^27^; briefly, each well was filled with a 1% (v/v) solution of poly(ethyleneimine) (PEI, Sigma-Aldrich 181978) in ddH_2_O for 10 minutes, 0.1% (v/v) glutaraldehyde (Sigma-Aldrich G6257) in ddH_2_0 for 30 minutes, then rinsed twice with ddH_2_O. The cell and pregel solution were then added to fill each well. The device and molds were incubated for 45 minutes at 37°C. The molds were removed from the stretching device and clamps and placed in small petri dishes 60 mm in diameter with 8 mL of hASC media to incubate for 7 days at 37°C and 5% CO_2_ with media changes every 48 hours. After plasma cleaning, all steps were performed aseptically in a cell culture hood.

**Figure 1.**
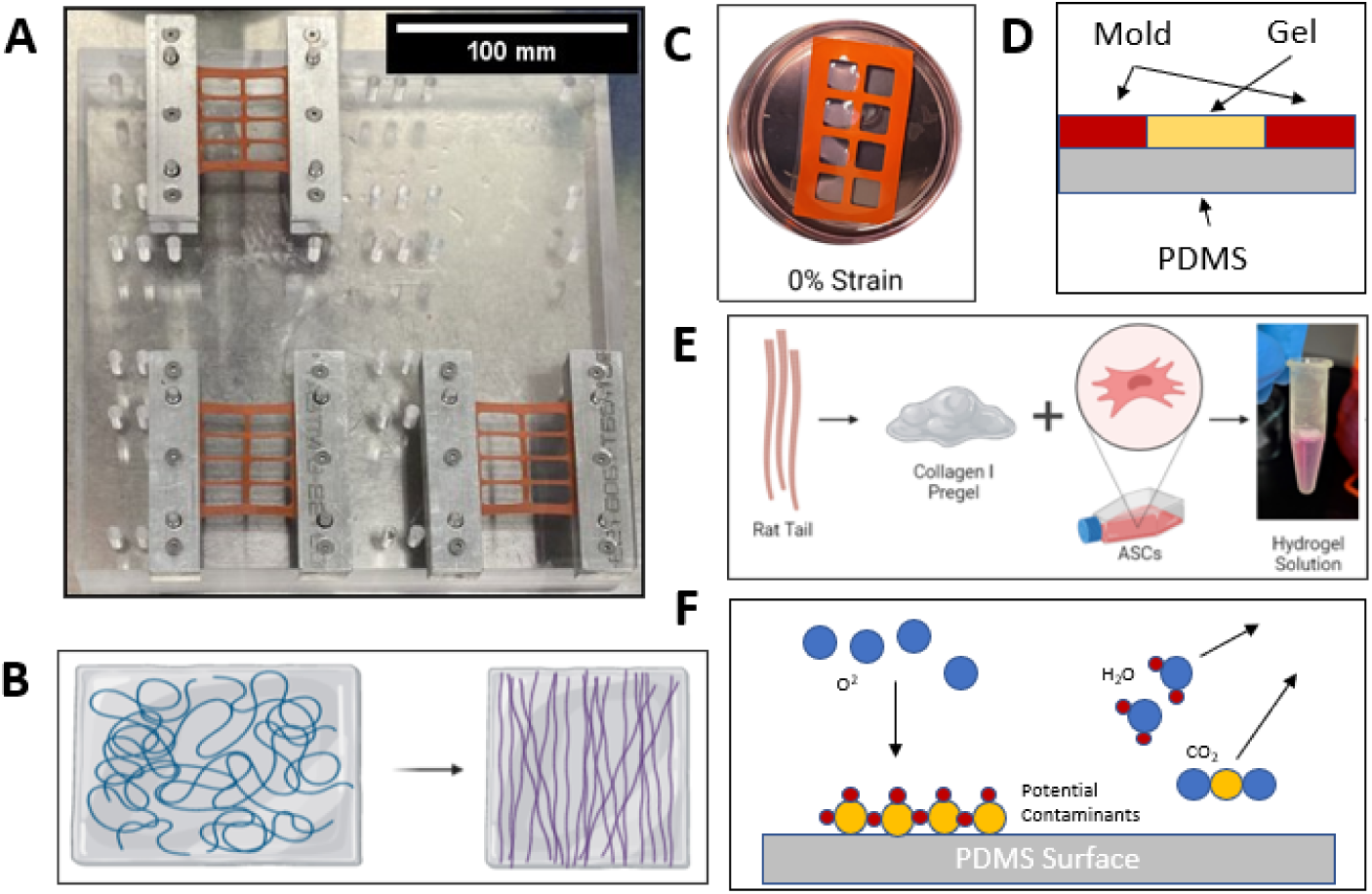
**A.** Stretching device comprising acrylic base, steel clamps, and silicone molds. Scale bar = 100 mm **B.** Schematic showing the relaxation of the pre-stretched molds leading to aligned fibers. **C.** Silicone mold and gels in petri dish with media. **D.** Cross-section of silicone mold. **E.** Hydrogel creation process. **F.** Depiction of plasma cleaner sterilization and sealing methods. Figures B, C, and E created with BioRender.com.

### Quantitative Polarized Light Imaging (QPLI)

Quantitative Polarized Light Imaging (QPLI) was performed on a previously established QPLI microscope^28^. Briefly, circularly polarized light is transmitted through a rotating linear polarizer driven by a stepper motor to generate linearly polarized light. The light is then focused with a condenser lens, transmitted through the sample, and collected with a 4x (0.13 NA) objective. Using a fixed circular analyzer and camera after the objective, 10 images (2056 x 2056 pixels, 1.4 µm/pixel) were collected in 18° increments of the rotating linear polarizer, and the oscillation in intensity at each pixel was used to calculate a pixelwise fiber orientation and phase retardation^28^. For this study, acellular stretched (S) and nonstretched (NS) gels with 4 different collagen I concentrations were placed on a microscope slide, covered with a coverslip, and imaged on the system. Due to the size of the gels, multiple images were collected and stitched together to create a full field of field for each gel, resulting in complete maps of fiber orientation and phase retardation. Following stitching of the images, masks were then generated by outlining the gel in the stitched image, and then applying a phase retardation threshold of 1°. To quantify fiber organization, an overall directional variance value, which is a measure of fiber organization ranging between 0 (anisotropic) and 1 (isotropic), was calculated from all fiber orientations within the generated mask^28^.

### Confocal Reflectance Microscopy

Acellular stretched and nonstretched hydrogels in 3 mg/mL and 6 mg/mL concentrations were created and fixed in 3.7% formaldehyde (Sigma-Aldrich 252549) in PBS at room temperature for 1 hour. The gels were then rinsed and stored in PBS until imaging at 640 nm with an Olympus IX83 confocal microscope (20x objective magnification, 1.8x digital zoom, NA 0.80) with a Z-step of 2 µm, starting from the ‘top’ surface and extending into the gel 250 slices or 500 µm. The resulting Z-stack was analyzed in ImageJ using the OrientationJ plugin. Evaluation within OrientationJ is based on the gradient structure tensor in an area within the region of interest^30^. The coherence of each sample was determined and used as a measure of alignment for the sample. Coherence values range from 0 to 1 and indicate orientation of image features, with 1 being anisotropic and 0 being entirely unaligned.

### Immunofluorescence Imaging

After 7 days of incubation, the gels were rinsed with PBS in their molds then fixed in 3.7% formaldehyde in PBS for 1 hour. The gels were rinsed once with PBS for 5 minutes then transferred to a 24 well plate. Each well was rinsed with PBS 2 more times for 5 minutes each. The gels were then rinsed with 0.1% triton X-100 (Sigma-Aldrich 93443-100ML) in PBS 2 times for 7 minutes. The gels were then incubated at room temperature with 1% bovine serum albumin (BSA, Sigma-Aldrich A7906-50G) in PBS for 1 hour. The gels were then allowed to sit overnight at 4°C with the primary antibody solution diluted in 1% BSA. The gels were then rinsed 3 times with 0.1% Tween 20 (Sigma-Aldrich P9416-100ML) in PBS for 5 minutes. Gels were then protected from light and incubated for 1 hour at room temperature with a secondary antibody solution in 1% BSA. The gels were then rinsed twice for 5 minutes with 0.1% Tween 20 and once with PBS before being stored in fresh PBS. Antibodies were purchased from Thermo Fisher unless otherwise stated, and their dilutions are listed as follows: DAPI (1:2500, D1306), Phalloidin 546 (1:500, A22283), Phalloidin 488 (1:500, A12379), anti-rabbit Laminin (1:200, PA1-16730), anti-mouse Fibronectin (1:200, MA5-11981), anti-mouse Yes-associated Protein (1:500, Santa Cruz Biotechnology, sc-101199), anti-mouse Neurofilament (1:500, Developmental Studies Hybridoma Bank, RT97), Alexa Fluor goat anti-mouse 488 (1:500, A11029), Alexa Fluor 488 goat anti-rabbit (1:500, A11008), Alexa Fluor goat anti-mouse 568 (1:500, A11031), Alexa Fluor goat anti-rabbit 568 (1:500, A11011), Alexa Fluor goat anti-mouse 647 (1:500, A21235), and Alexa Fluor goat anti-rabbit 647 (1:500, A21244). Fluorescence imaging was performed with an Olympus IX83 confocal microscope (20x objective magnification, NA 0.80) with a Z-step of 2 µm, starting from the ‘top’ surface and extending into the gel 250 slices or 500 µm.

### Live/Dead Assay

To determine cell viability in the scaffolds, 50 µL of Live/Dead solution (Fisher Scientific R37601) was added to each gel in the mold for 15 minutes. The gels were then suspended in clear Dulbecco’s Modified Eagle Medium (DMEM, Fisher Scientific 12-800-017) until imaging. Each gel was removed from its mold and placed on a microscope slide for imaging with an Olympus IX83 confocal microscope (20x objective magnification, NA 0.80) with a Z-step of 5 µm, starting from the ‘top’ surface and extending into the gel 100 slices or 500 µm. The percentage of live and dead cells was calculated with a custom MATLAB code quantifying the area occupied by live and dead cells. Analysis was performed on the compressed Z-stack at max intensity.

### Luminex

Serum-free ASC media were added to gels and collected after 24 hours to analyze hASC secretome. The conditioned media were briefly spun to remove cell debris, followed by tenfold concentration using Amicon Ultra-4 Centrifugal Filter (3 kDa MWCO, Millipore Sigma UFC800396). Custom-designed 4-Plex Luminex kits (R&D Systems) were used to quantify beta-nerve growth factor (βNGF), neurotrophin-3 (NT-3), glial derived neurotrophic factor (GDNF), and brain derived neurotrophic factor (BDNF) as well as hepatocyte growth factor (HGF), basic fibroblast growth factor (FGF-2), angiopoetin-1 (Ang-1), and interleukin-8 (IL-8) levels in the conditioned media. Resulting values were normalized respective to double-stranded DNA (dsDNA) content. DNA concentration was determined using DNeasy Blood and Tissue Kit (Qiagen 69504) to isolate dsDNA and Quantifluor dsDNA System (Promega E2670) to quantify DNA content.

### YAP Staining Analysis

Quantification of immunofluorescent images stained against YAP was performed by creating a Z-stack of slices 10-30 in ImageJ with constant minimum and maximum values for both YAP and DAPI intensity. These images were exported to MATLAB for analysis with a custom code determining the intensity of staining in the nuclear and cytoplasmic regions of the cells.

### Mechanical Testing

Acellular and cellular 1 million (M) cell/mL, 3 mg/mL hydrogels were used for rheologic testing after 7 days of incubation. A Discovery Hybrid Rheometer (DHR-2) equipped with 25 mm compression plate (TA Instruments, New Castle, DE) was used to test the viscoelastic properties of collagen gels. A plate gap of 1 mm was set before the gels were compressed a total distance of 995 μm at a rate of 31.5 μm/s. Data was collected in Trios software provided by TA Instruments and exported to excel for analysis. The Young’s Moduli were determined from the linear portion of the graphs.

### DRG Harvest and Dissociation

All animal work was approved by the Institutional Animal Care and Use Committee (IACUC) of the University of Arkansas (protocol numbers 23019 and 20054). Male Sprague Dawley rats (3-4 weeks) weighing 35-49 g were purchased from Envigo and cared for by staff of the South Campus Animal Facility (SCAF) in accordance with the IACUC Standards and the Animal Welfare Act. Rats had access to 12-hour light/dark cycles and standard food and water *ad libitum*. Rats were euthanized in ordinance with the American Veterinary Medical Association guidelines via carbon dioxide asphyxiation.

#### Harvest Methods

Immediately following euthanasia, the rat was transferred to a sterile hood and placed on an absorbent mat for dissection and DRG harvest. The rat was then sprayed with 70% ethanol, and electric clippers were used to shave the dorsal side of the animal. The skin was removed from the dorsal muscle and fascia with surgical scissors. Bone cutters were then used to sever the spinal column in the cervical and lumbar regions then to isolate the spinal column by cutting on either side of the of the spine from the cervical to lumbar dislocations. Following removal of the spinal column and trimming of any extra muscle, the spinal cord was removed via hydraulic extrusion with a 3mL syringe and PBS. Surgical scissors were then used to cut through the dorsal and thoracic sides of the spinal column, allowing for the lateral halves to be separated. The DRGs were visualized and removed carefully with sterile forceps, cleaned and trimmed, and then collected in hibernate media (Thermo Fisher, A1247501) on ice until use.

#### Dissociation methods

Dissociation and coculture of the DRGs are adapted from a previously established method^31^. First, the DRGs were collected in a microtube with 0.1% trypsin (Thermo Fisher, 15400054) and 1 mg/mL collagenase (Millipore Sigma, 10103578001) in PBS and allowed to incubate at 37°C for 50 minutes, agitating the microtube every 10 minutes. The microtube was then centrifuged at 300g for 5 minutes. The supernatant was subsequently removed and replaced with a suspension of 0.1% trypsin in PBS and allowed to incubate at 37°C for 10 minutes. After incubation, neurobasal media supplemented with 1% Pen/Strep (Thermo Fisher, 15-140-122) and 2% B27 (Thermo Fisher, A3582801) were added to the microtube at a 3:1 ratio (media: PBS), and the microtube was centrifuged again. The final supernatant was removed and replaced with fresh supplemented neurobasal media. The partially dissociated DRGs were resuspended in the solution and kept on ice until coculture.

### Neurite Outgrowth Model and Quantification

#### Coculture methods

Acellular and cellular scaffolds were incubated for 7 days in ADSC media before neuron coculture. Immediately prior to seeding, media were removed from the hydrogels. A second sterile silicone isolator was placed atop the original, raising the outer wall of the mold. A 50 mL suspension of neuron media containing one intact DRG was then added to each individual hydrogel and contained within the added isolator. The DRGs were moved to the center of the hydrogel if needed. The DRGs were allowed to incubate without additional media for 12 hours at 37°C to assist in anchoring of the DRG to the scaffold^31^. After 12 hours, 6 mL of supplemented neuron media were then added to the hydrogels, insuring no DRGs are displaced in the process, and incubated for another 36 hours at 37°C until fixation and staining. Immunofluorescence staining of the hydrogel cocultures was performed with anti-neurofilament antibodies for neurite visualization and phalloidin for ASCs.

#### Quantification

Overall orientation of DRG outgrowth was performed on a Z-stack of the entire Neurofilament channel in ImageJ with the Distribution tool in the OrientationJ plugin, which provides an orientation degree for each non-zero pixel. Histogram values were binned in 10-degree increments. Quantification of neurites was performed in FIJI using the Sholl analysis tool in the SNT plugin with a step size of 100 µm. The center of the DRG was used as the starting point, and neurites were quantified from 100 – 1400 µm.

### Statistical Analysis

Statistical analysis was performed in GraphPad Prism 9.5.1 using unpaired t-tests, one-way, two-way, and three-way ANOVAs with Tukey’s post-hoc analysis. Statistical significance was determined at p<0.05.

## Results and Discussion

### Physical Properties

#### Fiber alignment analysis

We assessed collagen fiber alignment using QPLI and confocal reflectance. QPLI is a popular imaging method used to determine orientation of birefringent fibers within tissues and samples. To optimize our hydrogel composition for subsequent cell culture, QPLI was performed on stretched and nonstretched samples in collagen hydrogel concentrations of 1, 2, 3, and 6 mg/mL to examine the collagen fiber alignment (n=4 per group). The 1 mg/mL and 2 mg/mL hydrogel samples failed to hold their shape and alignment after being released from the stretching device, and thus had no significant change in directional variance resulting from stretching. Directional variance measured with QPLI revealed significant decrease in variance (increase in alignment) after stretching in the 3 mg/mL and 6 mg/mL samples (Figure 2a). Both 3 and 6 mg/mL samples maintained their shape and structure after being released from the stretching device.

**Figure 2.**
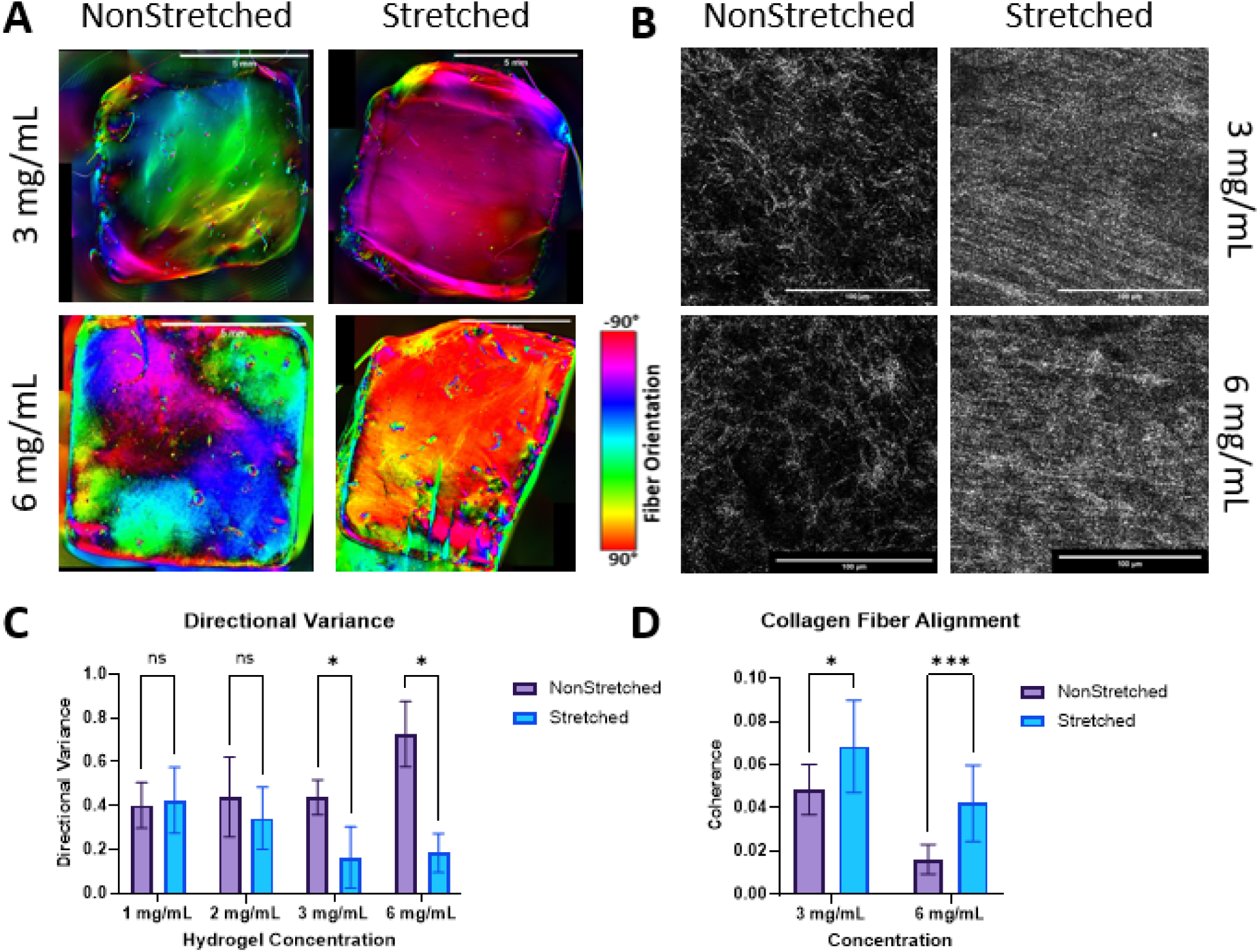
**A.** QPLI microscope images depicting fiber orientation in scaffolds after 1 day of culture, red is vertical fibers and cyan is horizontal. Scale bar = 5 mm. **B.** Confocal reflectance images visualizing collagen I fibers in acellular hydrogels after 2 days of culture. Scale bar = 100 µm. **C.** Graph showing the directional variance of samples with and without stretching. Analysis performed with multiple t-tests, *p < 0.05 for n=4/group. **D.** Graph showing the coherence (uniformity) for each sample group. Analysis performed with multiple t-tests, *p < 0.05, ****p < 0.001 for n=12/group.

To support our findings from QPLI, we performed confocal reflectance imaging on both stretched and nonstretched samples in 3 and 6 mg/mL concentrations to visualize individual collagen fiber organization and structure within the hydrogels. Imaging and analysis revealed significant increases in collagen fiber organization and alignment in both the 3 mg/mL and 6 mg/mL stretched samples compared to nonstretched controls (Figure 2b). Reflectance imaging of the collagen fibers resulted in organization that can be visualized at a macro-scale but is composed of many disconnected pixels that make analysis with pixel-based tools challenging. Due to this disconnection, the objective values for coherence of all reflectance samples, while indicating increased alignment, were lower than those seen in full fiber (actin) analysis. Similar results indicating an increase in alignment were seen with Sobel and Fourier analyses performed in MATLAB.

Through QPLI and confocal reflectance imaging, we were able to prove a significant increase in fiber alignment with the use of our stretching device. This result aligns with other studies achieving hydrogel fiber alignment through the usage of uniaxial stretch and compression^32,33^. Many of these studies utilize a one-piece, flexible PDMS mold, that can be pre-stretched or pre-compressed in their device, for the hydrogel to then be cast in and relaxed from. Unlike many of these studies, we have employed a two-part mold, comprising a silicone isolator and silicone sheet backer. This allows for easier removal of the hydrogels and their usage as a scaffold, as opposed to only an *in vitro* testbed.

### ASC Response

#### Cell alignment

After 7 days of incubation, 3 and 6 mg/mL hydrogels were stained with phalloidin to assess cell alignment within the hydrogels (Figure 3a). Image analysis revealed a significant increase in alignment in the stretched samples compared to the control in all cell densities and collagen concentrations tested (Figure 3b-c). There was an increase in cell alignment in the 6 mg/mL 1M samples compared to the same density in 3 mg/mL, but we did not see any significant difference in the 2M samples (Figure 3d).

**Figure 3.**
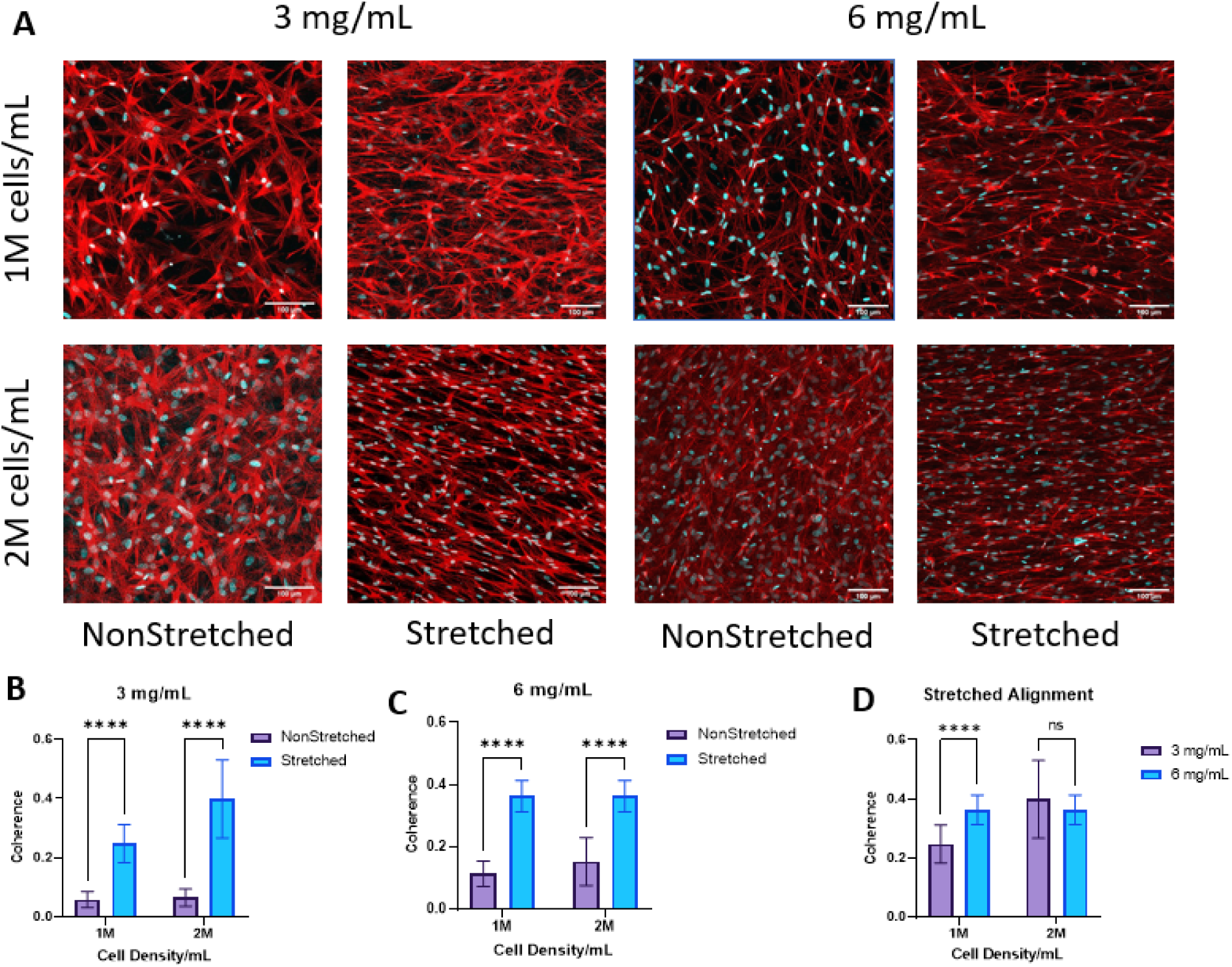
**A.** Images of cells in the bulk of the hydrogel after 1 week of culture, stained with phalloidin (red) to visualize f-actin and the cell body and with DAPI (blue) to visualize cell nuclei. Scale bar = 100 µm **B, C.** Graph depicting the coherence (alignment) of actin fibers within (**B**) 3 mg/mL and (**C**) 6 mg/mL samples. **D.** Graph showing the coherence of actin fibers within stretched scaffolds for 3 and 6 mg/mL and 1 M and 2 M/mL. Analysis performed with multiple t-tests. ****p < 0.0001 for n=12/group.

To confirm that cell alignment was coherent with collagen fiber alignment, we performed confocal reflectance imaging along with immunofluorescence to visualize cells and fibers together (Supplementary 1). Upon qualitative visualization of these results, we determined that cells were aligned uniformly along the direction of collagen fibers. This result is also in agreement with previous studies who have found co-alignment of encapsulated cells with scaffold fibers^32^. Not only can fiber alignment affect cell behavior and alignment, but this organization can provide structural guidance for outgrowing nerves to follow.

#### Cell Viability

After demonstration of collagen fiber alignment within the hydrogels, hASCs were added to the pregel before thermal gelation to allow for alignment of the embedded cells. To assess cell viability within the hydrogels, a live/dead assay was performed after 7 days of culture (Figure 4a). The assay revealed a significantly higher cell viability in the 3 mg/mL hydrogels compared to the 6 mg/mL samples (Figure 4c). Within each group, there was no difference in viability relating to cell density or stretched vs. control samples (Figure 4d-e). This is in agreement with published literature, with studies investigating the effects of both cell density and substrate alignment seeing no influence on viability from either factor^34^. So, while fiber alignment may influence cell morphology and behavior, it does not positively or negatively influence cell survival. For the above reasons, we decided to continue with only the 3 mg/mL collagen concentration group for further optimization and characterization. Because of the absence of significant alignment advantage and the decrease in cell viability in the 6 mg/mL samples, we decided against continuing with the 6 mg/mL concentration in this study.

**Figure 4.**
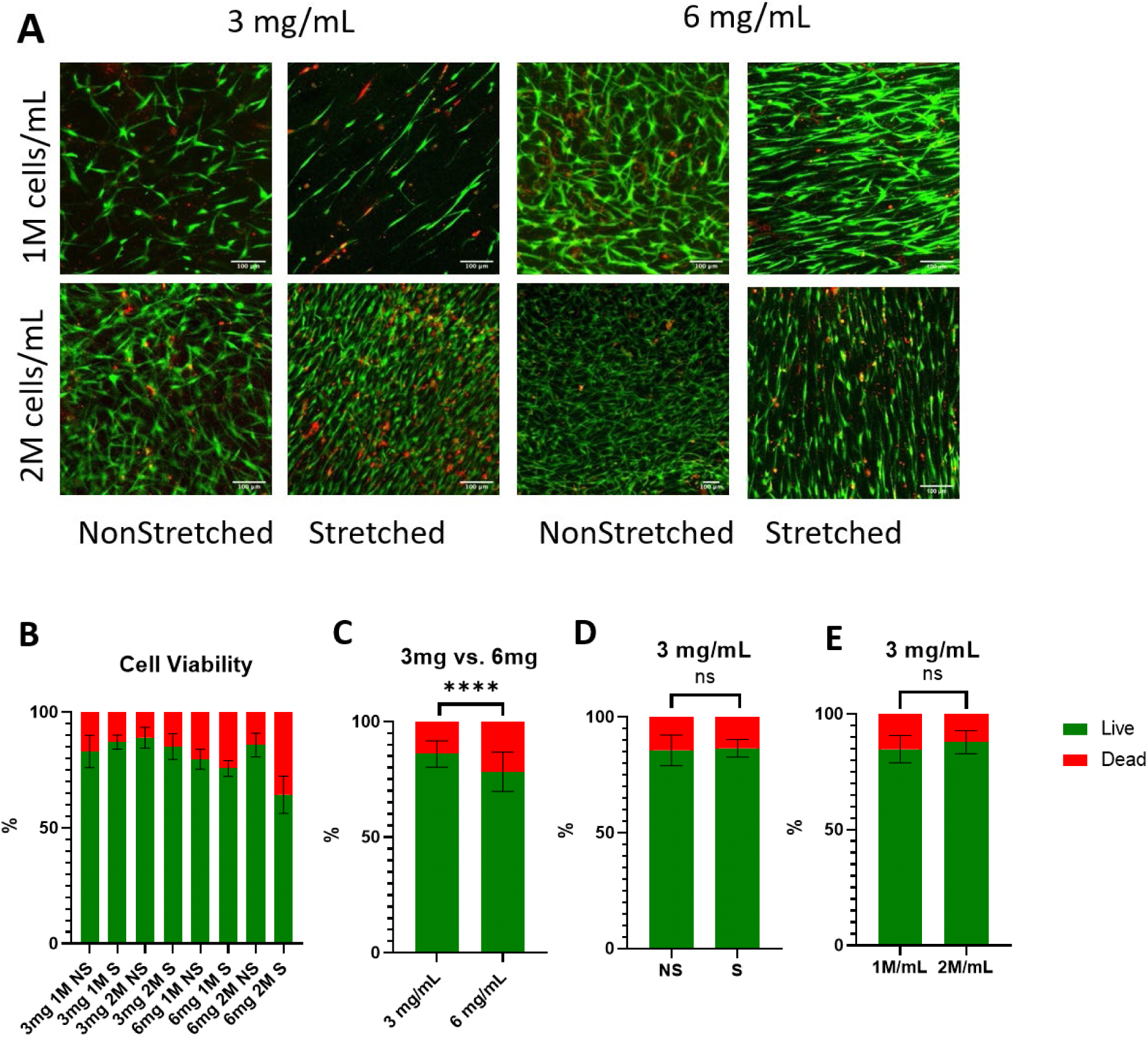
**A.** Images showing live (green) and dead (red) cells within the scaffolds after 1 week of culture. Scale bar = 100 µm **B.** Graph showing the percentage of live and dead cells within the scaffolds. Analysis performed with a one-way ANOVA and post-hoc Tukey’s test. **C.** Graph showing the percentage of live and dead cells in all 3 and 6 mg/mL samples. Analysis performed with t-test. ****p < 0.0001 for n=28-34/group. **D, E.** Graph showing the percentage of live and dead cells in 3 mg/mL samples, comparing (**D**) nonstretched (NS) and stretched (S) samples, and (**E**) 1 M/mL and 2 M/mL samples. Analysis performed with t-test. ns = no significance.

### ECM deposition and Cell Secretome

#### Cell Secretome

Conditioned media collected from 7-day hydrogel culture after 24 hours were analyzed with custom 4-plex Luminex kits (neurotrophic – NT-3, βNGF, GDNF, and BDNF; angiogenic - HGF, FGF-2, Angiopoetin-1, and IL-8) for hASC secretome in 3 mg/mL hydrogels with 1 M and 2 M/mL cell densities. NT-3 secretome was excluded from the results due to absence of expression in Luminex results. The 1 M/mL hydrogels showed significant increase in all analytes in the stretched samples compared to nonstretched for both neurotrophic and angiogenic factors (Figure 5). The 2 M/mL samples showed an increase in all angiogenic analytes as well as BDNF and βNGF in the stretched samples compared to the control (Figure 5).

**Figure 5.**
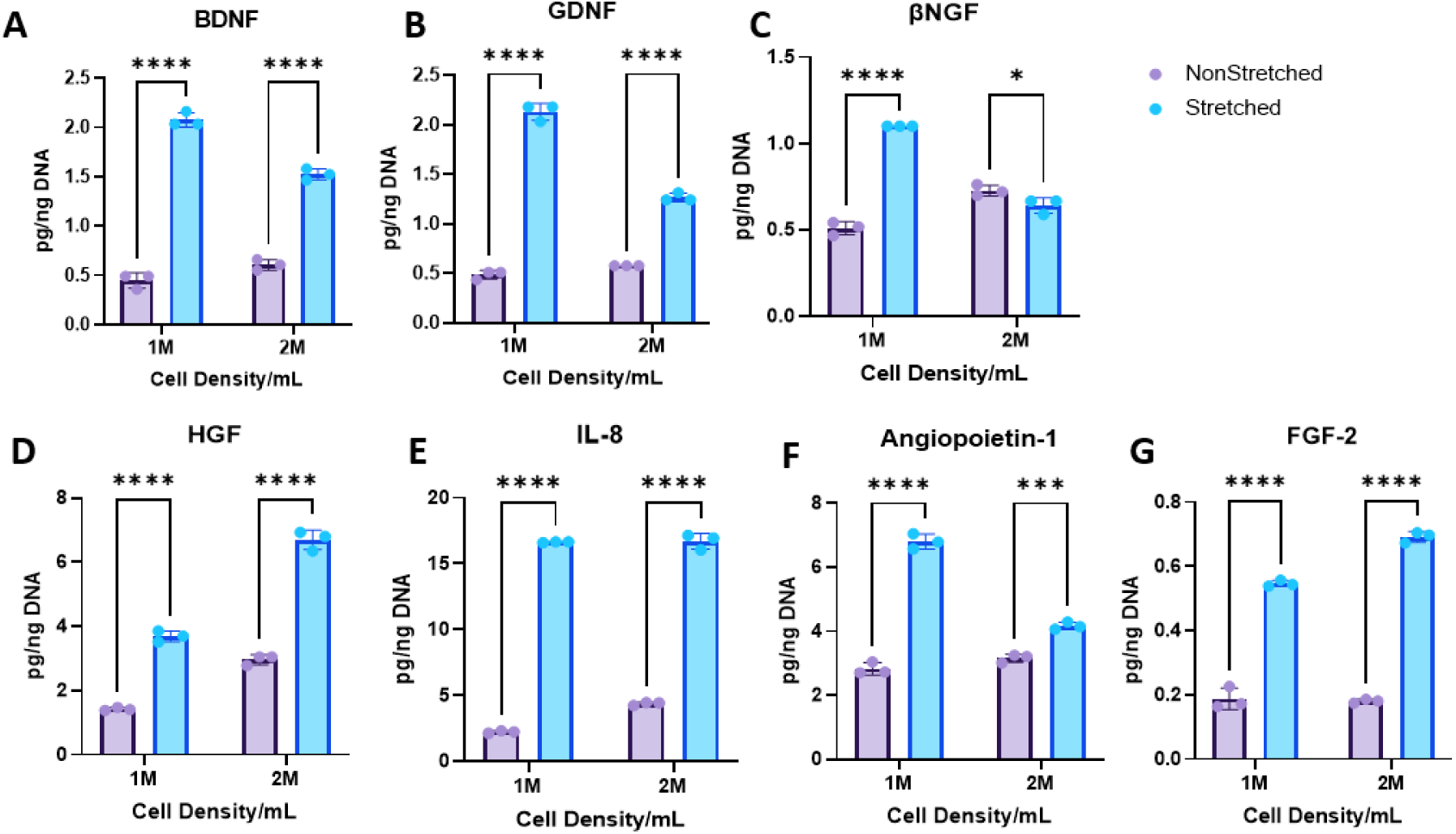
Graphs depicting results from Luminex analysis normalized to DNA content performed on conditioned media from 3 mg/mL scaffolds over 24 hours after 1 week of culture. Neurotrophins: **A.** Brain derived neurotrophic factor (BDNF) **B.** Glial derived neurotrophic factor (GDNF) **C.** Beta nerve growth factor (βNGF). Angiogenic factors: **D.** Hepatocyte growth factor (HGF) **E.** Interleukin 8 (IL-8) **F.** Angiopoietin-1 **G.** Fibroblast growth factor (FGF-2). Analysis performed with multiple t-tests. *p < 0.05, ***p < 0.001, ****p < 0.0001 for n=3/group.

Growth factors have been proven to be vital for translatable success with implantable peripheral nerve repair scaffolds^35^. Several studies have investigated the secretome response to cell culture in aligned substrates, and many saw increases in various immunomodulatory and angiogenic factors in MSCs and ASCs^36,37^. Neurotrophic factors, such as those mentioned above, have been proven to increase neurite outgrowth and functional outcomes in peripheral nerve repair applications ^38,39^, but no studies have investigated the influence of fiber alignment on MSC or ASC neurotrophic secretome. As previously mentioned, fiber alignment may influence neuro-directed stem cell differentiation, which may be positively influencing cell secretome.

In peripheral nerve repair, BDNF, GDNF, and NGF are well documented in literature and act to encourage proliferation, survival, and expression of peripheral neurons^38,39^. In addition to the benefits provided by the neurotrophic factors, the secreted angiogenic factors benefit peripheral nerve repair. Basic fibroblast growth factor (FGF-2) has been well documented in wound repair and vascularization, but supplementation of FGF-2 also has been proven to increase functional and morphologic outcomes such as myelination and Schwann cell proliferation after peripheral nerve injury^40^. While IL-8 is associated with both anti- and pro-inflammatory mechanisms and occasionally linked to pain at the injury site, studies have shown that increased IL-8 helps to recruit immune cells vital in wound healing and acts as an angiogenic factor encouraging vascularization, and the presence of which may be indicative of ASC to Schwann cell differentiation^41–43^. In another study, inclusion of Angiopoietin-1 (Ang-1) was able to establish vascularization early in peripheral nerve repair and is one of the two main factors in vascularization (VEGF being the other)^44^. HGF has been shown in additional studies to be vital in myelination thickness and axonal regrowth. It also promotes the proliferation and migration of Schwann cells and increases expression of endogenous GDNF^45,46^.

Encapsulation of cells stands as an attractive alternative to direct growth factor inclusion in scaffolds or local injection. Previously, growth factors have been administered at the site of injury with some success^35,38^. However, those factors are typically exhausted or not retained at the site of injury. A construct with encapsulated growth factors can ensure localization to the site of injury, but the growth factors will still be exhausted over time. In cellular scaffolds, the embedded cells will continue to produce growth factors as long as the contained cells survive. Another advantage of encapsulating cells directly, compared to growth factor encapsulation, is that multiple physiologically relevant growth factors can be incorporated with more ease than in acellular scaffolds. Because there was no obvious advantage in continuing with the 2 M/mL sample group, we selected the 1 M/mL sample group for further characterization and experiments.

#### ECM Deposition

After 7 days of incubation, 3 mg/mL hydrogels in 1 M and 2 M/mL cell densities were stained with antibodies against fibronectin and laminin (Figure 6). Fibronectin and laminin were chosen given their demonstrated roles in promoting nerve repair. Fibronectin plays a critical role in cell adhesion, migration, and proliferation, while laminin is a major component of the basement membrane and has been shown to play a crucial role in neural development and myelination^47–49^ . In addition, both fibronectin and laminin are known to influence maturation and function of Schwann cells. Imaging and orientation analysis revealed a significant increase in alignment of both laminin and fibronectin in the stretched samples for both densities (Figures 6c, e). Intensity analysis normalized to cell count showed the stretched 1 M/mL sample had the greatest deposition of laminin compared to all other samples (Figure 6d). The stretched 1 M/mL sample also showed the most deposition of fibronectin, while only significant against the stretched 2 M/mL group (Figure 6b). Although not statistically significant against the NS 1 M/mL and NS 2 M/mL groups, the difference of means is considerable, and with a larger sample size the significance should improve enough that we are confident drawing conclusions from this analysis.

**Figure 6.**
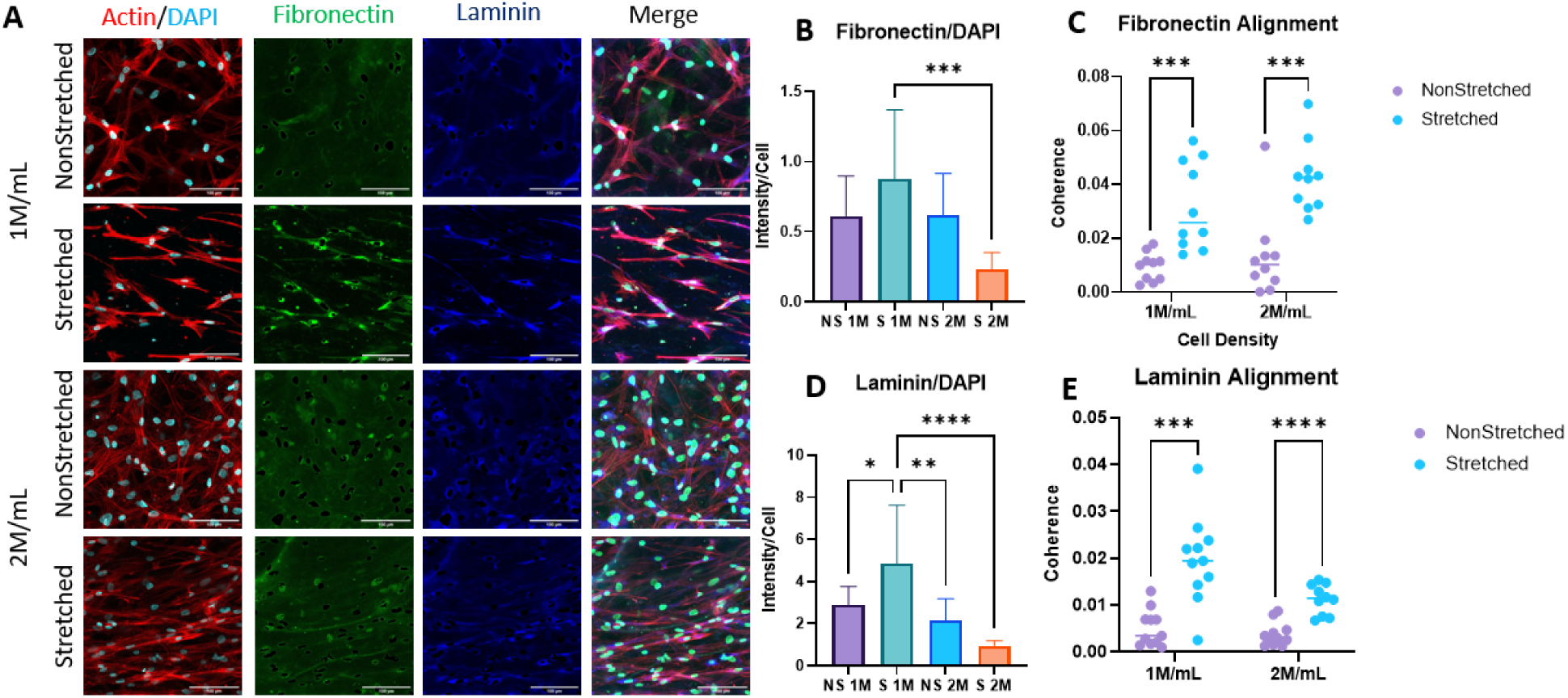
**A.** Immunofluorescence images depicting f-actin (red), fibronectin (green), laminin (blue), and DAPI (cyan) in 3 mg/mL scaffolds after 1 week of culture. Scale bar = 100 µm. **B.** Intensity normalized to cell count of fibronectin. **C.** Coherence (alignment) of fibronectin in scaffolds. **D.** Intensity normalized to cell count of laminin. **E.** Coherence of laminin in scaffolds. Intensity analysis performed with one-way ANOVA with post-hoc Tukey’s test. Orientation analysis performed with multiple t-tests. *p < 0.05, **p < 0.01, ***p < 0.001, ****p < 0.0001 for n=12/group.

While many studies have investigated benefits of included fibronectin and laminin in peripheral nerve constructs with success, their deposition by ASCs in response to substrate alignment has not been thoroughly explored^47–50^ . Additionally, fibronectin expression increases during the early stages of Schwann cell differentiation and is required for the development of myelin-forming Schwann cells^51^. Furthermore, laminin expression is upregulated during Schwann cell differentiation and is required for the proper alignment and spacing of Schwann cells during myelin formation^48^. Laminin also plays a role in regulating the expression of myelin genes in Schwann cells, which are necessary for the formation and maintenance of myelin sheaths^48^. ASCs possess the ability to differentiate into “Schwann-like” cells and may be doing so in response to cell and fiber alignment within the grafts. Spontaneous Schwannogenic differentiation of MSCs cultured in aligned substrates has been documented^52^, and while we have not stained against any Schwann cell markers in this study, ASC behavior suggests this as a possible explanation. Further investigation is needed to confirm this.

### Potential Mechanism (YAP)

To elucidate the mechanism driving ASCs’ increased ECM deposition and elevated expression of both neurotrophic and angiogenic secretome in stretched gels, we stained against Yes-associated Protein (YAP). The YAP and TAZ (Transcriptional co-activator with PDZ-binding motif) pathway is a signaling pathway involved in regulating cell growth, proliferation, and differentiation. YAP and TAZ are transcriptional co-activators that are downstream effectors of the Hippo pathway, which is a signaling pathway involved in tissue homeostasis. In the absence of YAP/TAZ activation, they are phosphorylated by the Hippo pathway kinases and sequestered in the cytoplasm, leading to their degradation^36,37,53^. Mechanical cues, such as substrate stiffness or fiber alignment, can activate the YAP/TAZ pathway and regulate its downstream effects. Specifically, when cells are subjected to mechanical forces that stretch or deform them, the resulting changes in cytoskeletal tension can lead to the activation and nuclear translocation of YAP/TAZ^36,37^.

The YAP/TAZ pathway has been implicated in elevated angiogenic secretome of MSCs and ASCs through mechanotransduction of both substrate stiffness and fiber alignment^36,37^. After staining against YAP, we saw a significant increase in YAP intensity in the stretched samples (Figures 7a-b). When we calculated the nuclear to cytoplasmic intensity ratio, however, we found that the nonstretched samples had a higher ratio with more proportional nuclear staining than the stretched group, which is usually found elevated when the pathway is triggered (Figure 7c). Nuclear staining was evident in both nonstretched and stretched samples.

**Figure 7.**
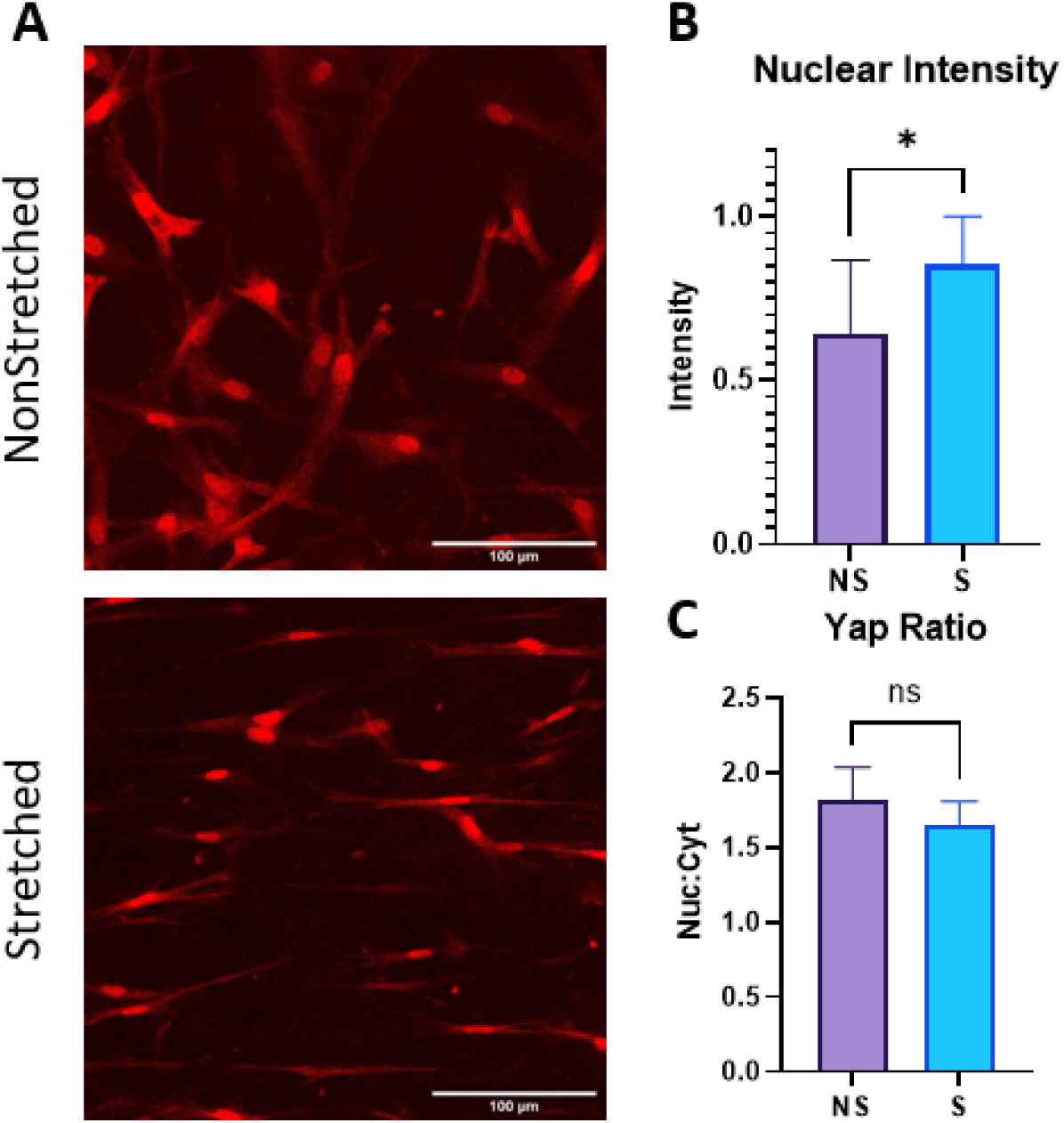
**A.** Images visualizing YAP (red) in 3 mg/mL 1 M/mL scaffolds after 1 week of culture. Scale bar = 100 µm. **B.** Nuclear intensity of YAP staining in scaffolds. **C.** Nuclear:cytoplasmic staining ratio for YAP. Analysis performed with t-tests. *p < 0.05 for n=12/group.

### Mechanical Testing

The stiffness of a hydrogel graft can not only influence embedded cell behavior, but native tissue behavior as well. Lower substrate stiffnesses (1-10kPa) influence cell differentiation and increase Schwann cell motility and proliferation^54^. Higher stiffnesses (>10kPa) have been shown to be beneficial for axon guidance in peripheral nerve repair grafts. Compression testing revealed Young’s moduli of all samples within the 1-10 kPa range, previously proven ideal for Schwann cell migration (Figure 8c)^54^. Further, the S 1 M/mL sample had a Young’s modulus of 6.5 ± 0.8 kPa, closest to a previously defined value of 7.45 kPa determined to be optimal for Schwann cell migration in a previously published study^55^.

**Figure 8.**
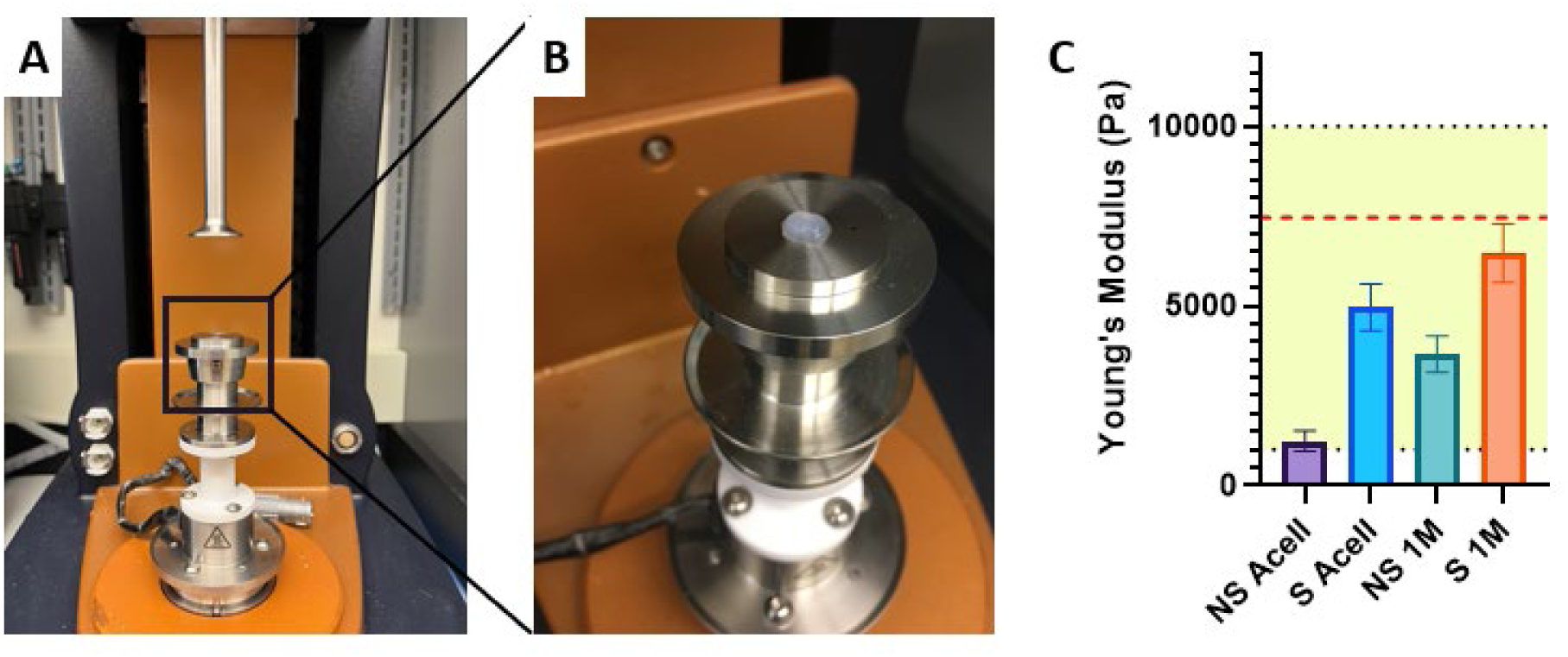
**A, B.** Images of compression testing setup. **C.** Graph visualizing Young’s modulus values for 3 mg/mL samples. Yellow shaded area representing 1-10 kPa and red dashed line representing 7.5 kPa target values. All groups significantly different at ****p < 0.0001 for n=22-30/group. Analysis performed with one-way ANOVA and post-hoc Tukey’s test.

### Neurite Outgrowth *in vitro*

To assess neurite outgrowth *in vitro*, partially dissociated, whole DRGs were seeded onto acellular and 1 M/mL nonstretched and stretched constructs (Figure 9a). This partial dissociation method, along with full dissociation and whole DRG culture, has been well documented in other studies^31,54,56,57^. Several studies have included NGF or other growth factors in their neuronal media; because we wanted to investigate the effects of cell alignment and their increased cell secretome on neurite production and extension, this was omitted in our media. Orientation analysis of the outgrowing neurites revealed the highest concentration of neurite alignment in stretched samples around 0 degrees, indicated by the peak (Figure 9b). The same analysis of nonstretched samples revealed no peak or localization of fiber orientation, suggesting a random, unstructured organization of the neurites. The -10-degree peak in the stretched sample group suggests primarily horizontal orientation of outgrowing neurites, beneficial when trying to bridge a longer nerve gap. The smaller peak around 80 degrees is perpendicular to the major orientation axis and represents the initial axon protrusion into the hydrogel. Often, axons growing into the hydrogel are unorganized close to the DRG, but as they grow farther from the source and into the aligned fiber network, they bend and eventually follow the orientation of cells and fibers.

**Figure 9.**
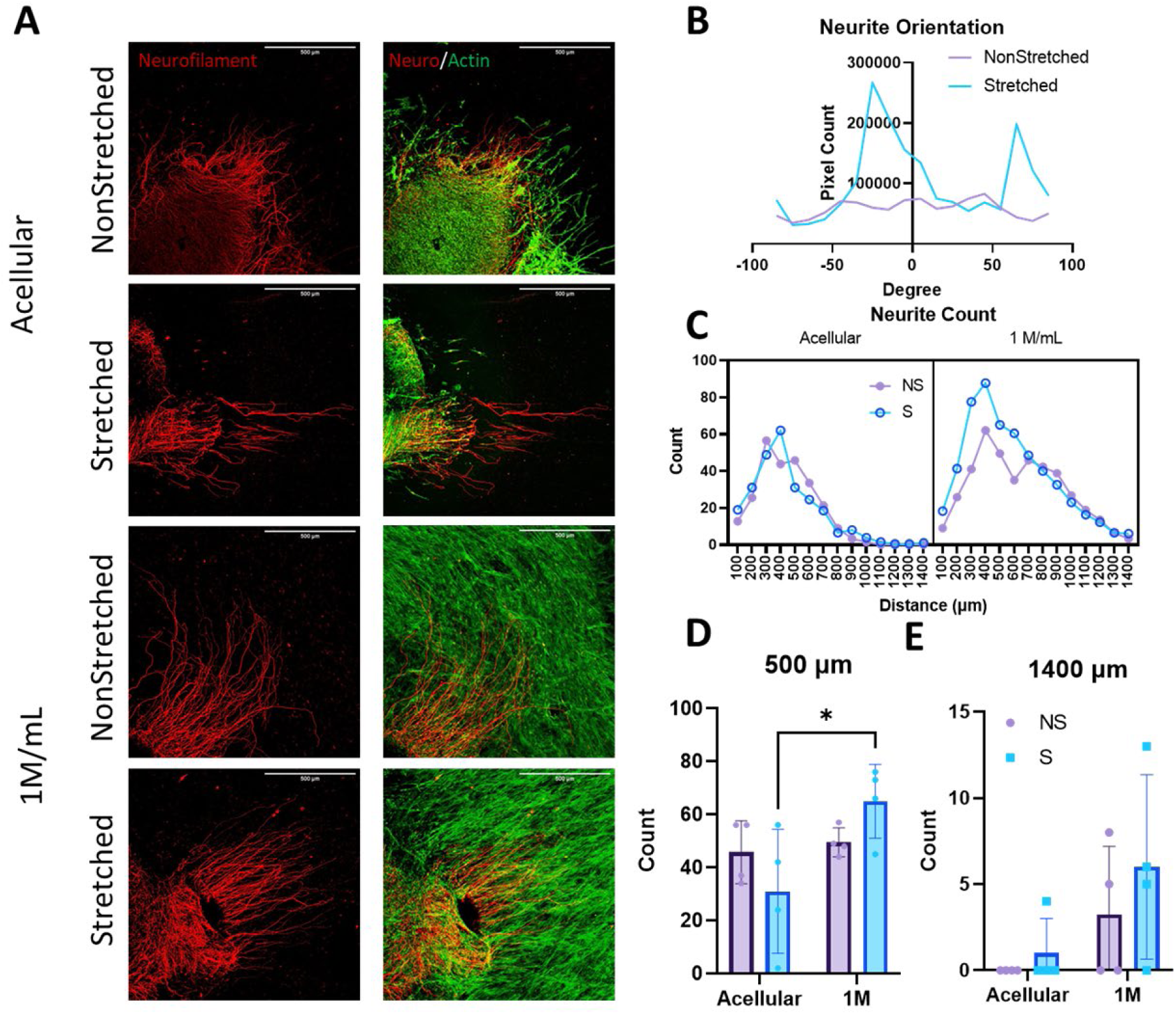
**A.** Immunofluorescent images of neurite outgrowth after 48 hours of co-culture on 3 mg/mL scaffolds visualizing neurofilament (red) and f-actin within cells (green). Scale bar = 500 µm **B.** Neurite orientation in 3 mg/mL 1 M/mL scaffolds, with values binned every 10 degrees. **C, D, E.** Neurite count over a certain distance: (**C**) 100-1400 µm, (**D**) 500 µm, (**E**) 1400 µm. Analysis for **C** performed with three-way ANOVA. Analysis for **D** and **E** performed with two-way ANOVA and post-hoc Tukey’s test. *p < 0.05 for n=4/group.

Neurite counts were performed with a Scholl analysis tool, providing a count of neurons reaching a distance; in this case, measurements were taken every 100 µm from 0 to 1400 µm (Supplementary 2). Sholl analyses function by creating concentric rings around a center point and measuring dendritic intersection with those rings. A count of neurites measured extending from the center of the DRG from 100 to 1400 µm shows the highest number of neurites reaching over both 500 and 1400 µm in the stretched 1 M/mL group (Figures 9 d, e). Only the count at 500 µm for the S 1 M against the S acellular sample proved to be significantly different, however this is not surprising with the small sample size (n=4/group). With increased sample size, we expect the significance to improve. Three-way ANOVA revealed significant sources of variation from both cell count and stretch (p<0.0001 and p=0.0391 respectively) over all count distances, further supporting the argument for the stretched 1M sample’s advantage. Additionally, the Sholl analysis is a radial assay, meaning all outgrowth from the center, not only in a certain direction is valued. If the distance was measured toward a specific endpoint along the axis of orientation (such as a nerve stump), we would expect to see further outgrowth in the stretched samples. These results are in alignment with other studies proving the advantage of anisotropic scaffold construction for peripheral nerve repair^31,57^.

## Conclusions and Future Directions

Here, we have demonstrated the fabrication and *in vitro* potential of our aligned collagen I scaffolds for use in peripheral nerve repair. Utilizing a silicone-based mold and pre-stretching with an in-house device, we demonstrated significant and effective collagen fiber and cell alignment, increased neurotrophic and angiogenic secreted factors, and increased deposition and alignment of ECM components important for effective and directed nerve repair, possibly due to the mechanosensitive YAP/TAZ pathway. Furthermore, our *in vitro* model of neurite outgrowth provided evidence supporting increased axonal regeneration and directional guidance governed by fiber and cell orientation within the scaffolds. Our unique approach to the construction of the stretching device and accompanying molds allows for removal of the cell-laden, anisotropic hydrogel from the device after fabrication and its subsequent function as a nerve repair scaffold *in vivo*. These scaffolds can be patient specific and personalized to the anatomy of the nerve gap or injury.

Future studies will focus on further optimization of the scaffolds and characterization of embedded cells for subsequent *in vivo* investigation. We have suggested that the ability of the encapsulated ASCs to differentiate into Schwann-like cells may be beneficial in longer term success of the scaffolds *in vivo*. We would like to investigate the environmental differentiation of these ASCs as well as the effects of intentional ASC differentiation on scaffold secretome, axon myelination, and overall neurite outgrowth within the scaffolds. Further investigation into potential mechanisms is needed, as well. While YAP pathway activation remains an attractive mechanistic explanation as indicated by literature and preliminary results, optimization of the staining and analysis protocols are needed, including further investigations into the YAP/TAZ pathway. Additionally, varying substrate stiffness would help to confirm mechanism activation. Finally, assessing cell/fiber alignment and neurite length at varying time points would be beneficial to our understanding of longer-term behavior and effects of these scaffolds.

## Acknowledgements

This work was funded by the National Institutes of Health through the award number P20GM139768 and PhRMA Foundation (YHS), Arkansas Biosciences Institute (YHS and J-WK), and Undergraduate Summer Research Grant Award from the University of Arkansas (DJ). Luminex assay was performed with help from the Almodovar lab (University of Arkansas Department of Chemical Engineering). We would like to thank Dr. Jeffrey Wolchok and Dr. Jorge Almodovar for their helpful discussions on the manuscript.

---

**Supplementary Figure 1.**
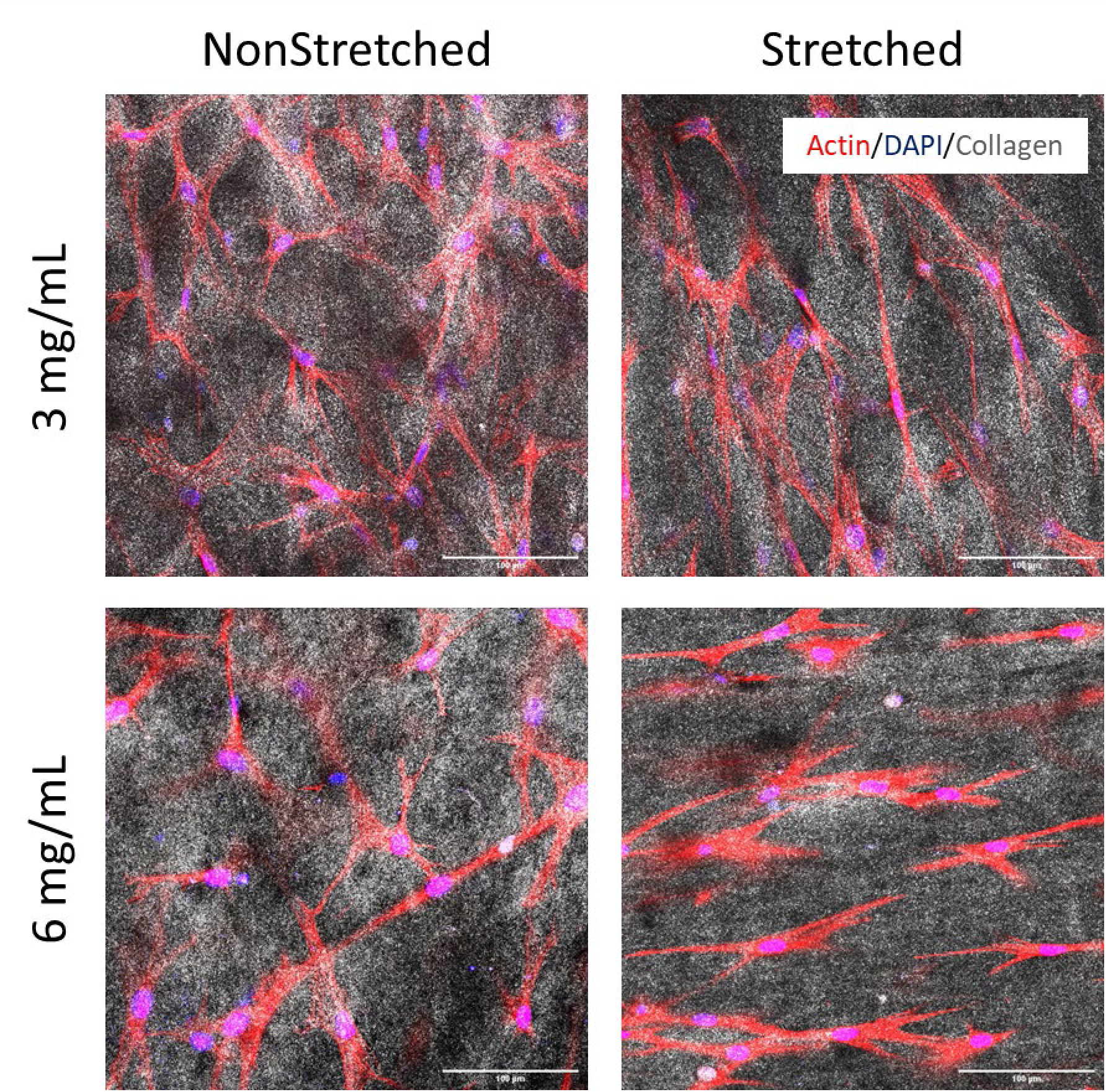
Images from confocal reflectance and immunofluorescence visualizing co-alignment of cells (red), nuclei (blue), and collagen fibers (gray) in 1 M/mL scaffolds after 1 week of culture. Scale bar = 100 µm

**Supplementary Figure 2.**
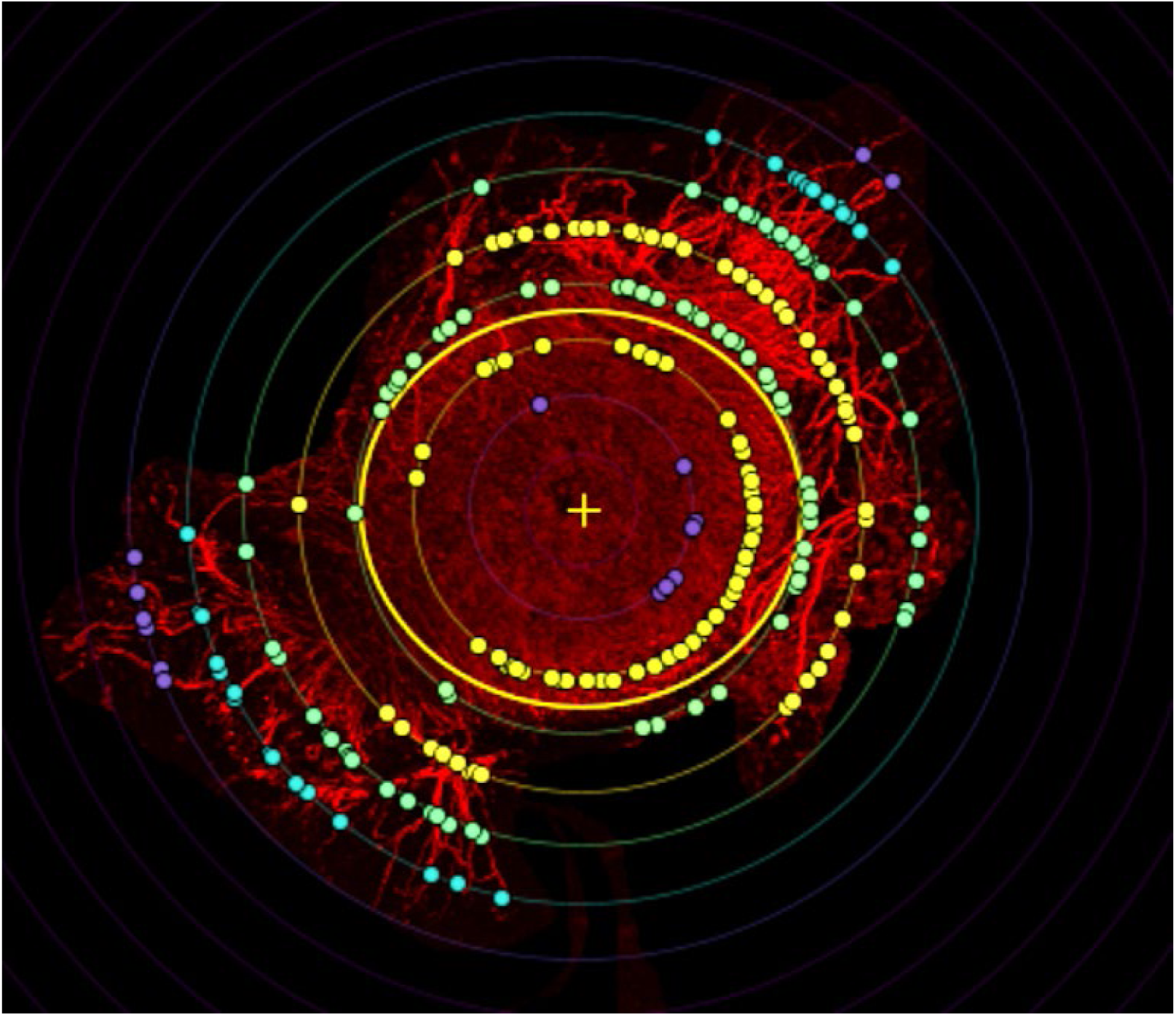
Image visualizing an example of Sholl analysis performed in FIJI. Yellow oval representing DRG body, yellow + indicating DRG center, used as starting point for outgrowth. Each concentric circle has color-coded points indicating neurite extension through the shell.

