## Supplementary figures and images for "Neuro-Regenerative Behavior of Adipose-Derived Stem Cells in Aligned Collagen I Hydrogels"

### Supplemental Figure 1

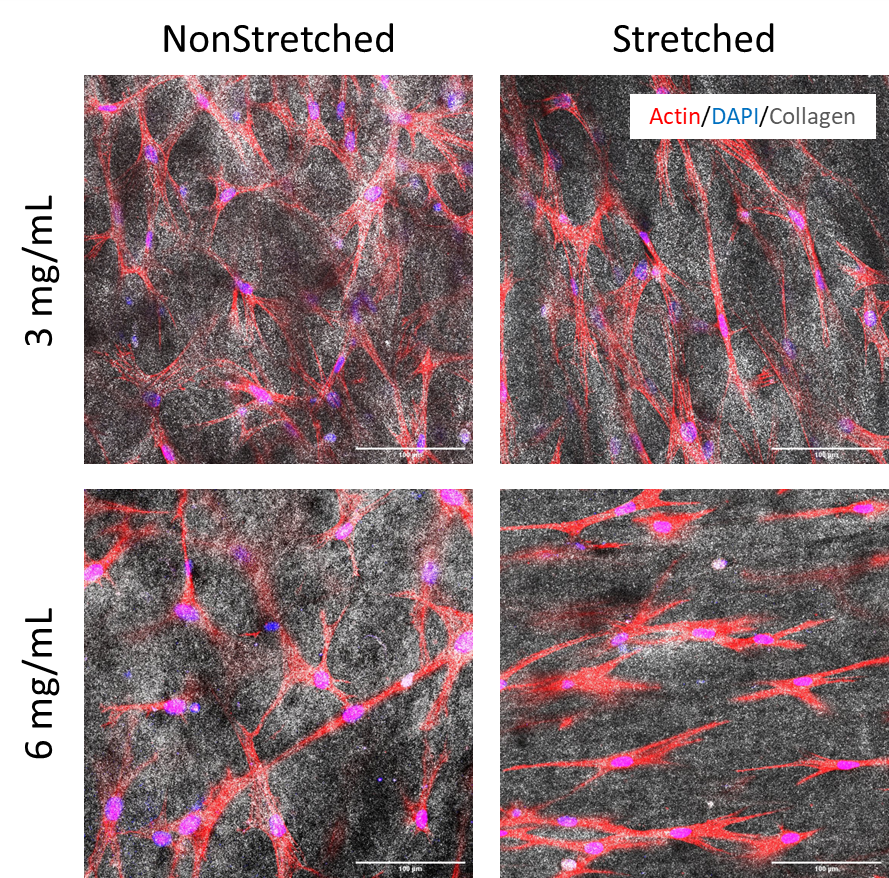

### Supplemental Figure 2

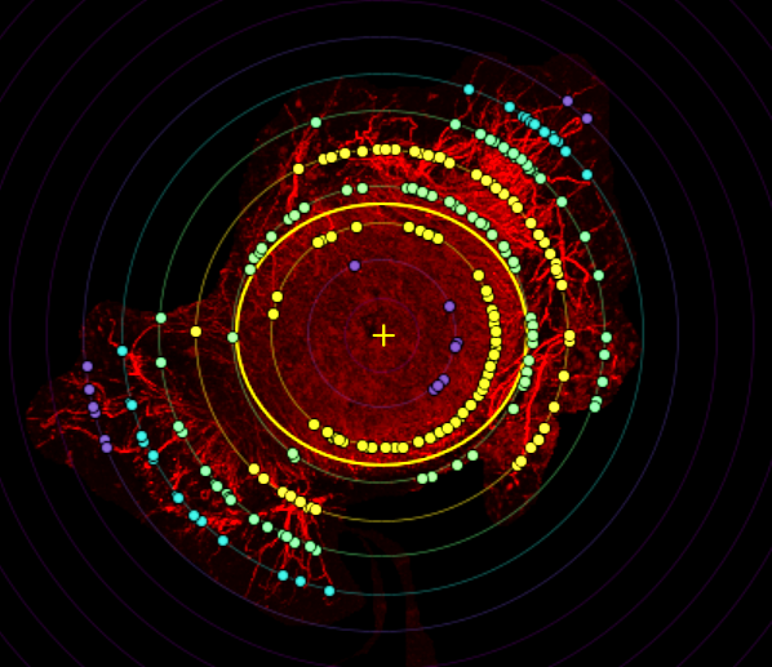
